# BindScreen: Protein-Centric Contrastive Learning for Sequence-Based Virtual Screening

**DOI:** 10.64898/2026.08.24.746801

**Authors:** Gabriel Bianchin de Oliveira, Fahad Saeed

## Abstract

Virtual screening ranks candidate molecules against a protein target. Sequence-based deep learning avoids docking’s structural requirements, but pair-based models need one forward pass per protein-molecule pair and scale poorly to large libraries. Dual-encoder contrastive models remove that bottleneck, yet standard CLIP training assumes a symmetric, one-to-one correspondence, whereas protein–molecule binding is asymmetric and many-to-many. We present *BindScreen*, a sequence-only dual-encoder screening model, and show that the decisive design choice is not the contrastive loss but how the batch is built. BindScreen combines a protein-centric batch construction and an asymmetric multi-positive InfoNCE loss. A factorial ablation separates the two contributions: the loss alone degrades performance under standard CLIP batching, the protein-centric batch alone recovers most of the gain, and the combination performs best. The effect is encoder-agnostic across eight protein language models spanning four architectural families. By decoupling protein count from molecule count per batch, BindScreen reaches higher validation BEDROC in 86 hours than standard CLIP reaches in 460 hours, and needs about seven times fewer forward passes to screen LIT-PCBA than pair-based models. The source code, pretrained checkpoints, and datasets are publicly available at https://github.com/pcdslab/BindScreen and https://huggingface.co/collections/SaeedLab/bindscreen.

## 1 Introduction

A fundamental step in drug discovery is identifying candidate molecules that target disease-related proteins, commonly referred to as drug-target interaction. However, experimental and laboratory based methods are notoriously time-consuming and expensive, with estimates indicating that developing a single marketable drug requires years of effort and billions of US$ in funding [1]. Virtual screening has emerged as a powerful computational technique to identify promising candidates for experimental validation by ranking large molecular libraries containing millions of compounds compared against a target protein. By reducing the number of compounds that need to be tested in the laboratory, virtual screening significantly cuts both time and cost [2].

Existing virtual screening methods rely on molecular docking, simulating protein-molecule interactions from three-dimensional structures of both the target and the compound [3–5]. While useful, these methods are computationally expensive and require 3D structural data that may not always be available - even with advances in structure prediction [6–8]. Deep learning methods have emerged as a powerful alternative, and can be divided into sequence-based approaches, which use amino acid sequences and SMILES strings [9–11], and structure-based approaches, which also use 3D information [12–15]. Consequently, sequence-based approaches offer a more broadly applicable alternative.

To fully leverage sequence data, recent research has shifted toward frameworks that map distinct biological modalities into a shared embedding space. Contrastive learning offers a way around the pairwise bottleneck: by learning a space in which binding proteins and molecules lie close together and non-binding ones far apart, each modality can be encoded independently, and screening reduces to a similarity search rather than one forward pass per pair. CLIP [16] demonstrated that this approach scales to large datasets, and it has inspired several adaptations for virtual screening. DrugCLIP [13] recasts screening as a retrieval task by aligning representations of protein pockets and molecules, achieving competitive performance against docking, and DrugHash [14] extends it with binary hash codes to reduce memory costs for large-scale screening. These methods, however, depend on three-dimensional pocket structures and, similarly to standard CLIP, assume a symmetric, one-to-one correspondence between the two modalities. Moreover, standard CLIP batching samples protein-molecule pairs at random, which accidentally penalizes valid biological pairings by treating them as negatives, limits the contrastive signal available per protein, and imposes substantial memory costs when training large sequence encoders. Two gaps thus motivate our approach, namely the reliance on 3D structure and the mismatch between CLIP’s symmetric batching and the many-to-many nature of binding.

In this work, we design and develop *BindScreen*, a sequence-based virtual screening method, built on a dual-encoder contrastive architecture, trained on 583,960 protein-molecule pairs spanning 1,492 proteins and 394,190 molecules curated from ChEMBL. For the external LIT-PCBA evaluation, we additionally re-filter this set to remove proteins similar to LIT-PCBA targets, preventing data leakage. BindScreen addresses the limitations of standard CLIP training through two design choices, an asymmetric multi-positive InfoNCE loss [17] that reflects the protein-centric nature of virtual screening, and a protein-centric batch construction strategy that increases the density of contrastive signals while reducing memory requirements. We compared our proposed model by conducting a systematic evaluation across 8 protein language models and 3 molecular language model variants, and assessed generalization on the LIT-PCBA [18] dataset against both sequence-based and structure-based methods. Our experiments demonstrate that the protein-centric batch construction improves EF@0.5 by up to 48% over standard CLIP training across all eight protein encoders, reaching a higher validation BEDROC in 86 hours than standard CLIP reaches in 460. On LIT-PCBA, BindScreen improves early enrichment over the retrained sequence-based baselines while requiring roughly seven times fewer forward passes than pair-based methods at inference.

The main contributions of this work can be summarized as follows:

- A factorial study that isolates batch construction from the loss function and identifies batch composition, not the contrastive objective, as the primary driver of the gains (up to 48% higher EF@0.5 over standard CLIP), with the asymmetric multi-positive loss degrading the performance under standard CLIP batching and becoming beneficial only when combined with the protein-centric batch.
- BindScreen, a sequence-only dual-encoder screening model whose protein-centric batch construction decouples the number of proteins from the number of molecules per batch, shifting the memory cost onto the smaller molecular encoder.
- A systematic evaluation showing that the effect is encoder-agnostic, with the protein-centric batch improving early-enrichment metrics across all 8 protein language models spanning four architectural families, frozen projection heads outperforming joint finetuning, and substructure-level pre-training proving most effective within the MolDeBERTa family.
- An analysis of training and inference efficiency, with the protein-centric strategy reaching a higher validation BEDROC in 86 hours than standard CLIP achieves in 460 hours, requiring roughly 7 times fewer inference forward passes than pair-based methods, and yielding molecular embeddings reusable across targets.
- A held-out LIT-PCBA evaluation using a sequence-clustered split, together with an analysis of why strong internal enrichment does not transfer to out-of-distribution targets.

## 2 Methods

In this section, we present the BindScreen method, the datasets, the baseline methods, and the evaluation metrics.

### 2.1 BindScreen Model

BindScreen uses a dual-encoder architecture that independently encodes each modality using specialized language models, as illustrated in Figure 1. The protein branch processes amino acid sequences through a protein language model, while the molecule branch extracts SMILES representations using a molecular language model. In both branches, the final-layer token representations are mean-pooled to produce a single embedding per input.

**Figure 1:**
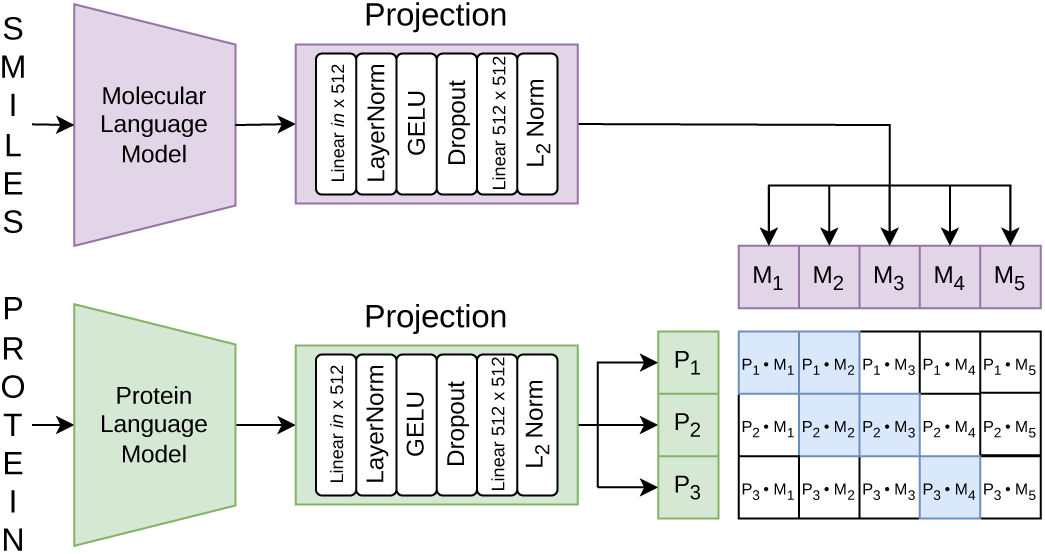
BindScreen pipeline. Protein sequences and SMILES strings are independently encoded by a protein language model and a molecular language model, respectively. Both representations are passed through projection heads to produce normalized embeddings. The batch contains *P* proteins (rows) and *M* molecules (columns), where blue cells indicate known binding pairs (positives) and black cells indicate in-batch negatives.

To align the two embedding spaces, each representation is passed through a projection head comprising a linear layer with 512 units, layer normalization, GELU activation, dropout with probability equal to 0.1, and a second linear layer with 512 units, followed by *L*_2_ normalization so that each embedding lies on the unit hypersphere. The projection dimension of 512 was chosen as a balance between the molecular encoder output (768 dimensions) and the protein encoder outputs (≥1024 dimensions), avoiding both excessive compression and unnecessary expansion.

To assess the impact of encoder architecture on virtual screening performance, we evaluate 8 protein language models spanning four distinct paradigms. Encoder-only models are represented by ESM C [19], ESM1b [20], ESM2 T36 [7], and ProtBERT [21], encoder-decoder models by Ankh 3 XL [22] and ProtT5 [21], autoregressive generative models by ProGen2 [23], and convolutional models by CARP 640M [24]. On the molecular side, we evaluate three variants of the MolDeBERTa [25] base model. Each variant was trained with a different objective, i.e., the MLM variant reconstructs masked SMILES tokens, the MTR variant predicts physicochemical and biological properties, and the MLC variant predicts the presence of molecular substructures.

Virtual screening is a protein-centric task, where given a query protein the goal is to rank candidate molecules by binding likelihood, and a single protein may bind multiple molecules. The loss should therefore support multiple positives per anchor. Standard InfoNCE handles neither of these properties, as it assumes a single positive per anchor and is applied symmetrically in both directions. We therefore adopt an asymmetric multi-positive InfoNCE loss [17], applied in the protein-to-molecule direction only. We restrict the loss to this direction because screening is inherently protein-anchored, ranking candidate molecules for a query protein. Equation 1 defines this objective, where *p_i_* and *m_j_* are the normalized projected embeddings of protein *i* and molecule *j*, *τ* is a learnable temperature parameter initialized to 0.1, *B* is the number of molecules in the batch, Q(*i*) is the set of known binding molecules for protein *i*, and *P* is the number of proteins in the batch.

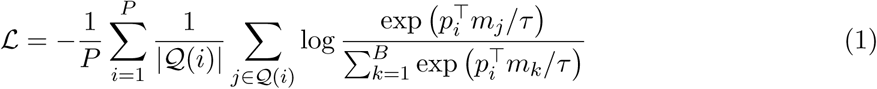

Beyond the loss function, the efficacy of contrastive learning depends heavily on batch composition. Standard CLIP training builds batches by randomly sampling protein-molecule pairs. This limits the contrastive signal available per step and, with large encoders, severely constrains the batch size, since jointly backpropagating through both architectures is memory-intensive. To overcome these limitations, we propose a protein-centric batch construction strategy. Given a target batch of *P* proteins and *B* molecules, we sample *P* proteins and assign *M* = ⌊*B/*(2*P* )⌋ known binding molecules to each, forming an initial set of explicit positive pairs. The remaining *B* − (*P* × *M* ) molecules are sampled at random from the training set and shared across all proteins in the batch. Crucially, every pair is checked against the ground-truth annotations after the batch is assembled, rather than being fixed by each molecule’s sampling origin. As a result, a randomly sampled molecule may become a positive for one or more proteins, and a molecule assigned to one protein may also bind another in the same batch. This exposes multiple positives per protein, naturally supports many-to-many interactions, and reduces false negative supervision while preserving molecular diversity. The number of proteins per batch *P* controls a trade-off, where a larger *P* increases the diversity of protein anchors but assigns fewer binders to each, whereas a smaller *P* concentrates more positives on fewer anchors. This process is illustrated in Figure 2.

**Figure 2:**
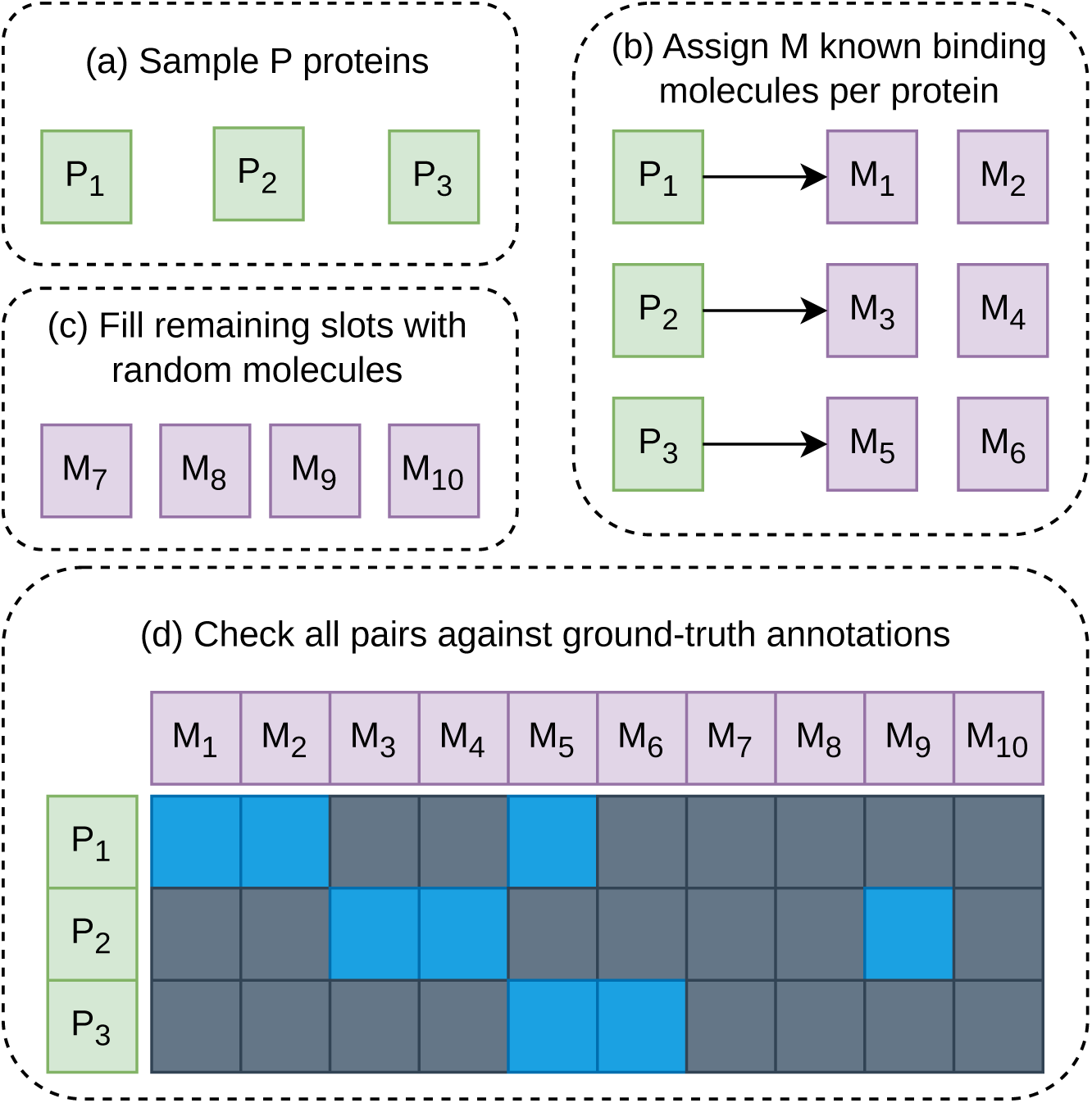
Protein-centric batch construction process. (a) *P* proteins are sampled to form the batch. (b) Each protein is assigned *M* known binding molecules, producing an initial set of explicit positive pairs. (c) The remaining *B* − (*P* × *M* ) slots are filled with molecules sampled randomly from the training set. (d) After the batch is assembled, every protein-molecule pair is checked against the ground-truth binding annotations. Blue cells indicate positive pairs, which may come from explicit assignment (e.g., *M*_1_ and *M*_2_ for *P*_1_), from a molecule assigned to one protein that is also a known binder of another (e.g., *M*_5_, assigned to *P*_3_, is also positive for *P*_1_), or from a randomly sampled molecule that matches a known binding annotation (e.g., *M*_9_ for *P*_2_). Gray cells indicate negatives within the batch.

### 2.2 Datasets

We built our training dataset from ChEMBL 36 [26], following established protocols in the literature for data collection and filtering [13, 27]. The query was restricted to binding assays targeting single proteins with small molecules, with activity measurements limited to Ki, Kd, IC50, and EC50 values reported in nM with an exact relation. Duplicate protein-molecule pairs were resolved by taking the median activity value, molecules with more than 100 heavy atoms were removed, and proteins with fewer than 30 ligands or fewer than 5 active compounds were excluded. All compounds with an activity value below 562,000 nM were labeled as active, following the threshold adopted by Wang et al. [27]. Our training set had a total of 583,960 pairs, spanning 1,492 proteins and 394,190 molecules.

Rather than splitting the dataset randomly, we clustered all protein sequences using MMseqs2 [28] with a minimum sequence identity of 0.4 and coverage of 0.8, and assigned clusters to splits in 70% for training, 15% for validation, and 15% for testing. This sequence-based split produces a more realistic evaluation setting, where the model is tested on proteins that are genuinely distinct from those seen during training. For the internal ChEMBL evaluation, each test protein serves as a query and is ranked against all molecules in the test split. Molecules annotated as binders of the query protein are treated as actives, and all remaining molecules as assumed negatives.

To compare our model against the literature, we evaluated BindScreen on LIT-PCBA [18], a widely used virtual screening dataset comprising 15 protein targets with experimentally validated active and inactive compounds. To prevent data leakage, we re-filtered the training and validation sets by removing any protein with sequence similarity to a LIT-PCBA target, using the same MMseqs2 parameters. Table 1 summarizes the statistics of all dataset splits used in this work.

**Table 1:** Dataset statistics for the ChEMBL and LIT-PCBA splits.

| Dataset | Split | Proteins | Molecules | Pairs |
| --- | --- | --- | --- | --- |
| ChEMBL | Train | 1,492 | 394,190 | 583,960 |
|  | Validation | 334 | 101,586 | 147,676 |
|  | Test | 326 | 105,048 | 130,281 |
| ChEMBL filtered | Train | 1,452 | 375,120 | 548,294 |
|  | Validation | 326 | 97,213 | 143,088 |
| LIT-PCBA | Test | 15 | 404,586 | 2,776,973 |

The batch-construction, encoder-selection, and finetuning experiments use the full ChEMBL splits, whereas the LIT-PCBA comparison uses the filtered splits. No filtered test split is required, as evaluation in that setting is performed on LIT-PCBA itself. Unlike the ChEMBL splits, which contain only active compounds, LIT-PCBA includes both active and inactive molecules for each target.

### 2.3 Baselines

We compare BindScreen against two groups of methods. The first group consists of sequence-based drug-target interaction methods, which we retrained on our dataset. These methods are trained on explicit positive and negative pairs, whereas BindScreen derives its negatives from in-batch contrast at a much larger scale. The comparison is therefore controlled for training data but not for the amount of negative supervision, a difference we return to in the discussion. The second group consists of structure-based virtual screening methods, for which we report published results as a high-level reference, since they were trained on different datasets and use 3D structural information unavailable to BindScreen.

For the sequence-based group, we evaluate three methods. DeepDTA [9] encodes protein sequences and SMILES strings independently using token embeddings followed by convolutional layers, concatenates the resulting representations, and passes them through a multi-layer perceptron for binding prediction. HyperAttentionDTI [11] also relies on two convolutional encoders, but uses an attention mechanism for representation pooling before concatenating both outputs and applying a classification layer. Finally, MolTrans [10] processes each input through a Transformer-based encoder and combines the resulting representations via an interaction module before classification. We restricted the sequence-based comparison to methods with publicly available implementations. For each baseline, we used the original authors’ implementation and default hyperparameters without further tuning, which may understate their performance.

For the structure-based group, we include both traditional docking methods (Glide-SP [3], Gnina [4], and Surflex [5]) and learning-based methods that rely on 3D protein structures (Planet [29], BigBind [12], DrugCLIP [13], DrugHash [14], and S2Drug [15]). We report their published results solely as a reference point to contextualize the performance of sequence-based approaches, without claiming direct comparability.

### 2.4 Evaluation Metrics

We evaluate BindScreen using metrics designed for virtual screening, where the goal is to rank active compounds as high as possible in a list of candidates.

The Enrichment Factor at a given fraction *α* measures how much more concentrated active compounds are in the top *α*% of the ranked list compared to a random selection [30]. Equation 2 defines EF*_α_*, where *n_α_*is the number of actives in the top *α*% of the ranked list, *N_α_* is the total number of compounds in the top *α*%, *A* is the total number of actives, and *N* is the total number of compounds. In the internal ChEMBL evaluation, *A* is the set of annotated binders of the query protein and *N* is the full test pool, with all remaining molecules treated as assumed negatives, while in the LIT-PCBA evaluation, actives and inactives are the experimentally validated labels provided with the benchmark. We report EF at 0.5%, 1.0%, and 5.0%, which reflect early enrichment performance.

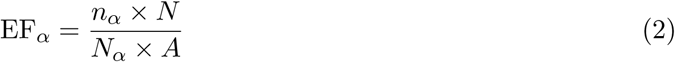

The Area Under the Receiver Operating Characteristic Curve (AUCROC) measures the overall ability of the model to separate actives from inactives across all possible thresholds. Although AUCROC is widely used, it treats all ranking positions equally and can be dominated by performance in the lower part of the ranked list, which is less relevant for virtual screening. To address this limitation, we also report the Boltzmann-Enhanced Discrimination of Receiver Operating Characteristic (BEDROC) [31] with *α* equal to 85, which assigns exponentially higher weights to compounds ranked at the top of the list. All metrics are computed per protein and averaged across the evaluation set.

### 2.5 Implementation Details

All BindScreen models were trained for 30 epochs on a single NVIDIA RTX A6000 GPU using the AdamW [32] optimizer. When training with unfrozen encoders, we used LoRA [33] to reduce memory requirements and a learning rate of 1 × 10^−5^, while in all experiments the projection heads used a learning rate of 1 × 10^−4^. The best model was selected based on BEDROC on the validation set. We fixed the total number of molecules per batch at *B* equal to 2,048, while the number of proteins *P* (and hence the assigned binders per protein *M* ) varied across configurations, as described in Section 3.

The sequence-based baseline methods require labeled pairs of active and inactive compounds for training. Since our ChEMBL dataset contains only active pairs, we generated an equal number of negative pairs by randomly sampling protein-molecule combinations not present in the dataset. This procedure was applied to both the training and validation sets. We used the original implementations of DeepDTA^1^, HyperAttentionDTI^2^, and MolTrans^3^, retrained on our dataset.

## 3 Results

In this section, we present the results of BindScreen across four experiments. We first evaluate the impact of the protein-centric batch construction strategy, then assess the choice of protein and molecular encoders, followed by an analysis of fine tuning strategies and training efficiency, and finally compare BindScreen against sequence-based and structure-based methods on LIT-PCBA. All results are reported on the hold-out test set.

### 3.1 Protein-Centric Batch Construction

First, we evaluate the impact of batch construction on virtual screening performance. In all experiments, both encoders are kept frozen and only the projection heads are trained. For each protein encoder, we swept over different configurations of *P* (proteins per batch) and *M* (known binding molecules per protein), keeping the total batch size fixed at 2,048. For each experiment, we selected the epoch that maximized BEDROC on the validation set. The complete results are provided in the Supplementary Material.

Table 2 compares the best protein-centric configuration with asymmetric loss against standard CLIP training with symmetric loss for each protein encoder, with MolDeBERTa MLC fixed as the molecular encoder. The protein-centric strategy consistently outperforms standard CLIP across all eight encoders, with gains on every early-enrichment metric and ties only on AUCROC for a single encoder, demonstrating that the improvement is not specific to a particular protein encoder architecture. The gains are especially pronounced in EF@0.5 and EF@1.0, which reflect early enrichment, the most relevant scenario in virtual screening practice.

**Table 2:** Comparison between standard CLIP and protein-centric batch construction for each protein encoder, using MolDeBERTa MLC as the molecular encoder. **Bold** indicates the best result for each metric within each encoder comparison.

| Protein Encoder | Batch | EF@0.5 | EF@1.0 | EF@5.0 | AUCROC | BEDROC |
| --- | --- | --- | --- | --- | --- | --- |
| Ankh 3 XL | Standard CLIP | 27.308 | 19.056 | 7.204 | 0.760 | 0.205 |
|  | Protein-Centric | **37.983** | **24.260** | **7.896** | **0.770** | **0.253** |
| CARP 640M | Standard CLIP | 28.597 | 19.271 | 7.474 | 0.761 | 0.207 |
|  | Protein-Centric | **39.568** | **24.871** | **8.112** | **0.773** | **0.258** |
| ESM C | Standard CLIP | 23.491 | 17.249 | 7.121 | 0.763 | 0.185 |
|  | Protein-Centric | **32.680** | **21.019** | **7.422** | **0.764** | **0.221** |
| ESM1b | Standard CLIP | 32.016 | 21.354 | 7.645 | 0.758 | 0.225 |
|  | Protein-Centric | **39.637** | **25.532** | **8.226** | **0.774** | **0.263** |
| ESM2 T36 | Standard CLIP | 30.961 | 21.158 | 7.736 | 0.767 | 0.224 |
|  | Protein-Centric | **39.615** | **25.520** | **8.626** | **0.797** | **0.267** |
| ProGen2 | Standard CLIP | 25.271 | 17.732 | 7.068 | 0.762 | 0.189 |
|  | Protein-Centric | **35.382** | **22.437** | **7.346** | **0.768** | **0.232** |
| ProtBERT | Standard CLIP | 22.551 | 16.626 | 7.029 | 0.759 | 0.179 |
|  | Protein-Centric | **33.332** | **21.420** | **7.226** | 0.759 | **0.222** |
| ProtT5 | Standard CLIP | 27.001 | 18.869 | 7.222 | 0.757 | 0.200 |
|  | Protein-Centric | **37.966** | **24.708** | **7.855** | **0.790** | **0.246** |

A key aspect of this comparison is that standard CLIP training operates on a 2,048 × 2,048 similarity matrix, while the protein-centric strategy computes a *P* × 2,048 matrix, where *P* ≪ 2,048. Despite requiring fewer operations, the protein-centric approach consistently achieves higher performance, suggesting that the quality of the contrastive signal, driven by protein diversity within the batch, matters more than the raw number of comparisons. This results in simultaneous gains in memory efficiency and model effectiveness.

The best configurations vary across encoders, suggesting that different protein representations benefit from different levels of protein diversity within the batch. Models such as ESM2 T36 perform best with moderate protein diversity, with 32 proteins per batch, while others such as CARP 640M and ProGen2 benefit from higher diversity, with 1,024 proteins per batch. ProGen2, the only autoregressive encoder, shows one of the largest gains, with EF@0.5 increasing from 25.3 to 35.4, suggesting that autoregressive models may particularly benefit from a richer contrastive signal during training for virtual screening. A plausible explanation is that the number of in-batch comparisons is *P* × *B* across protein-centric configurations, so what varies is not the amount of contrastive signal but its composition, namely the ratio of positives per anchor and the diversity of negatives seen by each protein. Encoders whose protein representations are less separated in embedding space appear to require greater protein diversity within the batch to produce informative gradients, consistent with ESM2 T36 favoring *P* = 32 while CARP 640M and ProGen2 favor *P* = 1,024.

To isolate the contribution of each proposed component, Table 3 presents a factorial ablation study using ESM2 T36 and MolDeBERTa MLC, independently varying the batch construction strategy and loss function. For the protein-centric configuration with symmetric loss, we used *P* equal to 32 and *M* equal to 32, matching the best configuration identified for ESM2 T36 in Table 2. Protein-centric batch construction alone consistently outperforms standard CLIP training, indicating that it is the primary driver of the observed performance gains. In contrast, applying the asymmetric loss under standard CLIP batch construction slightly degrades performance across all metrics, suggesting that the asymmetric objective alone is insufficient to improve virtual screening performance. When both components are combined, the model achieves the best overall performance, indicating that the asymmetric objective becomes most effective when coupled with the protein-centric batch construction. Together, these results suggest that the two components are complementary, i.e., the protein-centric batch construction provides the contrastive structure necessary for the asymmetric objective to operate effectively, while the asymmetric loss further improves the quality of the learned representations within this setting.

**Table 3:** Ablation of batch construction and loss function using ESM2 T36 and MolDeBERTa MLC. **Bold** indicates the best result per metric.

| Batch | Loss | EF@0.5 | EF@1.0 | EF@5.0 | AUCROC | BEDROC |
| --- | --- | --- | --- | --- | --- | --- |
| Standard CLIP | Symmetric | 30.961 | 21.158 | 7.736 | 0.767 | 0.224 |
| Standard CLIP | Asymmetric | 27.651 | 19.539 | 7.427 | 0.760 | 0.207 |
| Protein-centric | Symmetric | 35.411 | 23.384 | 8.126 | 0.780 | 0.244 |
| Protein-centric | Asymmetric | **39.615** | **25.520** | **8.626** | **0.797** | **0.267** |

### 3.2 Encoder Selection

Based on the results in Table 2, ESM2 T36 achieved the highest BEDROC among all evaluated protein encoders and was selected for the remaining experiments. Then, we evaluated the three MolDeBERTa variants to determine the most suitable molecular encoder, keeping ESM2 T36 and the protein-centric batch construction with 32 proteins fixed.

The results, presented in the Supplementary Material, show that MolDeBERTa MLC outperforms MLM and MTR in EF@0.5, EF@1.0, EF@5.0, and BEDROC, while MTR and MLM achieve slightly higher AUCROC. This suggests that substructure-based pre-training captures molecular features more relevant to binding than token reconstruction or property regression. The ability to recognize molecular substructures may provide a more direct inductive bias for virtual screening, where the presence of specific chemical groups often determines binding affinity.

### 3.3 Finetuning and Training Efficiency

After selecting ESM2 T36 and MolDeBERTa MLC as the best encoders, we evaluated the impact of fine tuning on model performance. We compare four configurations, both encoders frozen, full finetuning of both encoders, finetuning only MolDeBERTa, and finetuning only ESM2 T36, with the results presented in the Supplementary Material.

Fine tuning both encoders simultaneously degrades performance across all metrics compared to the frozen baseline, and finetuning MolDeBERTa alone produces a similar drop. Finetuning only ESM2 T36 performs comparably to the frozen configuration, with small and mixed differences across metrics and an identical BEDROC (0.267), our model selection metric. We therefore adopt the frozen configuration as our main model for its simplicity and lower training cost, and report the finetuned variant as an ablation. These results suggest that the projection heads are sufficient to align the two embedding spaces, and that updating the molecular encoder introduces noise that degrades the contrastive signal. A possible explanation is that the MolDeBERTa fine tuning process on the ChEMBL dataset pushes the representations away from their original generalization capacity. In contrast, finetuning ESM2 T36 alone does not degrade performance. We hypothesize that the protein sequence space is more constrained than the molecular space, allowing the encoder to adapt its representations without losing generalization.

Next, we compare our protein-centric strategy against standard CLIP training under the same setup, fine tuning only the ESM2 T36 encoder. Unlike the frozen setting, where the method use the 2,048-molecule batch, finetuning the encoder makes the joint gradient computation memory-bound and forces standard CLIP to a much smaller batch, which changes the nature of the comparison. Standard CLIP training limits the batch size to 32 proteins and 32 molecules due to memory constraints, resulting in only 1,024 effective protein-molecule comparisons per batch and requiring approximately 92 hours per epoch. In contrast, our protein-centric strategy operates with batches containing 32 proteins and 2,048 molecules, yielding 65,536 effective protein-molecule comparisons per batch while requiring approximately 2.8 hours per epoch. In wall-clock terms, the protein-centric strategy completes all 30 epochs in approximately 86 hours and reaches a higher validation BEDROC, whereas standard CLIP requires more than 460 hours to complete only five epochs, as shown in Figure 3. This corresponds to a roughly 32× reduction in per-epoch training time, driven by decoupling the number of proteins from the number of molecules in the batch.

**Figure 3:**
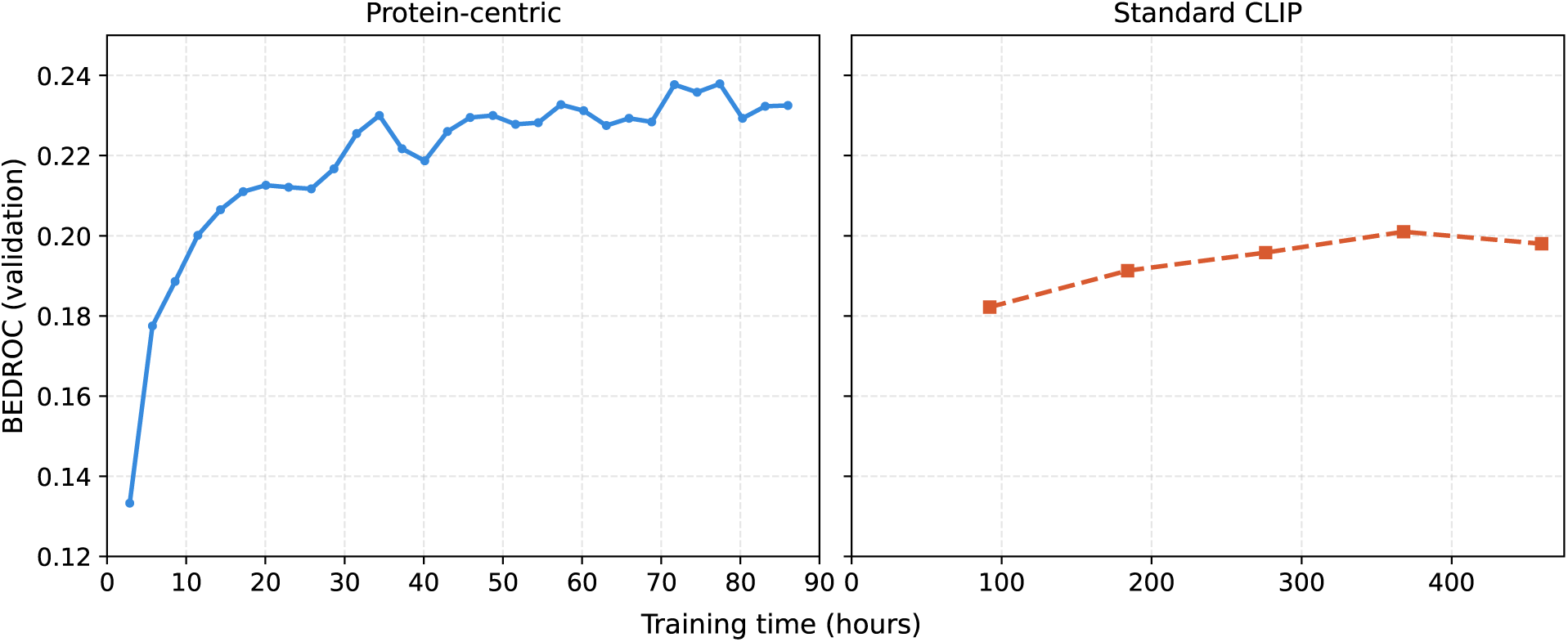
Validation BEDROC over training time for protein-centric batch construction (left) and standard CLIP (right).

### 3.4 Comparison with the Literature

Finally, we evaluated BindScreen on LIT-PCBA, reporting our main frozen configuration (BindScreen F) alongside the finetuned-only ESM2 T36 variant (BindScreen FT) as an ablation. Table 4 presents the results alongside the sequence-based baselines retrained on our dataset and structure-based methods reported in the literature.

**Table 4:** Comparison of BindScreen against sequence-based (top) and structure-based (bottom) methods on LIT-PCBA. BindScreen F refers to the frozen configuration and BindScreen FT refers to the ESM2 T36 finetuned configuration. Methods marked with * report results from DrugCLIP paper, + report results from DrugHash paper, and - report results from S2Drug paper. **Bold** indicates the best result for each metric within each group.

| Method | EF@0.5 | EF@1.0 | EF@5.0 | AUCROC | BEDROC |
| --- | --- | --- | --- | --- | --- |
| BindScreen F (ours) | 3.02 | **2.62** | **1.46** | **0.537** | **0.034** |
| BindScreen FT (ours) | 3.41 | 2.42 | 1.19 | 0.532 | 0.030 |
| DeepDTA | 1.48 | 1.20 | 1.10 | 0.500 | 0.022 |
| HyperAttentionDTI | 2.45 | 1.91 | 1.39 | 0.530 | 0.027 |
| MolTrans | 1.53 | 1.25 | 1.70 | 0.531 | 0.026 |
| Surflex* | - | 2.50 | - | 0.515 | - |
| Glide-SP* | 3.17 | 3.41 | 2.01 | 0.531 | 0.040 |
| Gnina* | - | 4.63 | - | **0.609** | 0.054 |
| Planet* | 4.64 | 3.87 | 2.43 | 0.573 | - |
| BigBind* | - | 3.82 | - | 0.608 | - |
| DrugCLIP* | 8.56 | 5.51 | 2.27 | 0.572 | 0.062 |
| DrugHash<sup>+</sup> | 9.65 | 6.14 | 2.42 | 0.546 | 0.060 |
| S2Drug<sup>-</sup> | **11.44** | **7.38** | **2.97** | 0.582 | **0.087** |

Among the retrained sequence-based methods, BindScreen (frozen) achieves the best EF@0.5, EF@1.0, AUCROC, and BEDROC. On EF@5.0, MolTrans remains competitive (1.70 versus 1.46), indicating that our advantage concentrates in the early-enrichment regime. These early-enrichment gains suggest that the protein-centric contrastive approach learns more transferable representations than pair-based classification methods. Beyond predictive performance, BindScreen also offers a significant inference efficiency advantage over pair-based methods. While methods such as DeepDTA, HyperAttentionDTI, and MolTrans require one forward pass per protein-molecule pair, BindScreen encodes proteins and molecules independently and computes similarities via dot product. On LIT-PCBA, this reduces the number of forward passes from 2,776,973 (one per pair) to 404,601 (404,586 molecules plus 15 proteins), approximately 7 times fewer. Furthermore, molecular embeddings can be precomputed and reused across different target proteins, making BindScreen particularly efficient in large-scale screening scenarios where the same molecular library is screened against multiple targets.

Compared to structure-based methods, BindScreen achieves lower enrichment factor values than DrugCLIP, DrugHash, and S2Drug, which rely on 3D protein structures. Despite this gap, BindScreen’s frozen EF@0.5 (3.02) falls in the same range as the docking baseline Glide-SP (3.17), while remaining below it, using only sequence information and no 3D structure. This is a relevant operating point, since structure-based methods require 3D protein structures that are not always available, whereas BindScreen operates directly from amino acid sequences.

The absolute enrichment factor values on LIT-PCBA are substantially lower than those observed on the internal ChEMBL test set, which is expected. LIT-PCBA is a particularly challenging dataset, with highly imbalanced active-to-inactive ratios and only 15 proteins. Furthermore, our sequence-based split ensures that similar LIT-PCBA proteins are unseen during training, making the evaluation more stringent than random splits commonly used in the literature. These factors together explain the performance difference between the two evaluation settings.

## 4 Discussion

In this work, we presented BindScreen, a sequence-based virtual screening method built on a dual-encoder contrastive architecture. Its two design choices, an asymmetric multi-positive InfoNCE loss and a protein-centric batch construction, target the asymmetric, many-to-many nature of protein-molecule binding. A factorial ablation showed that the gains are driven primarily by the batch construction rather than the loss, and that the two components are complementary. The protein-centric strategy consistently outperformed standard CLIP across all eight protein encoders while improving memory and training efficiency, reaching a higher validation BEDROC in 86 hours than standard CLIP reached in 460. Among retrained sequence-based baselines, BindScreen achieved the strongest early enrichment, and at inference it required roughly seven times fewer forward passes than pair-based methods. On LIT-PCBA, its sequence-only enrichment was competitive with, though below, structure-based screening.

BindScreen has several limitations. Negatives are generated by random sampling, which may not capture the true distribution of inactive chemical space, and the retrained baselines use fewer negatives than our in-batch contrast, so the comparison is not controlled for negative supervision. The internal ChEMBL evaluation relies on assumed negatives, and the large gap between internal and external enrichment indicates that generalization to out-of-distribution targets remains an open challenge. Performance also varies considerably across protein encoders, showing that representation quality limits overall performance. Our results were obtained with a single seed, the three molecular encoders belong to the same MolDeBERTa family, and the external evaluation is restricted to the 15 targets of LIT-PCBA.

As future work, the independence of the two encoders allows the ranking signal to be probed directly by re-ranking with permuted protein embeddings, isolating target-specific from target-independent contributions. We also plan to evaluate additional molecular encoders and virtual screening datasets, and to incorporate predicted structural information into the protein representation.

## Acknowledgments

The authors thank the anonymous reviewers for their valuable suggestions. This research was supported by the NIGMS of the National Institutes of Health (NIH) under award number: R35GM153434. The content is solely the responsibility of the authors and does not necessarily represent the official views of the National Institutes of Health.

## Protein-Centric Batch Construction Sweep

Tables S1-S8 present the full sweep of protein-centric batch construction configurations for each protein encoder, using MolDeBERTa MLC as the molecular encoder. For each configuration, *P* denotes the number of proteins per batch and *M* the number of known binding molecules per protein, with the total batch size fixed at *B* = 2048. **Bold** indicates the best result for each metric within each encoder.

**Table S1:** Full sweep of protein-centric batch construction configurations for Ankh 3 XL.

| P | M | EF@0.5 | EF@1.0 | EF@5.0 | AUCROC | BEDROC |
| --- | --- | --- | --- | --- | --- | --- |
| 1 | 1024 | 33.897 | 22.567 | 7.814 | 0.790 | 0.228 |
| 2 | 512 | 32.959 | 21.887 | 7.850 | 0.790 | 0.227 |
| 4 | 256 | 35.298 | 23.069 | 8.053 | 0.795 | 0.239 |
| 8 | 128 | 35.390 | 23.574 | 8.308 | **0.801** | 0.243 |
| 16 | 64 | 35.392 | 23.720 | **8.401** | 0.800 | 0.249 |
| 32 | 32 | 36.120 | 23.970 | 8.239 | 0.785 | 0.249 |
| 64 | 16 | 35.503 | 23.457 | 8.051 | 0.789 | 0.245 |
| 128 | 8 | 36.047 | 23.336 | 7.996 | 0.778 | 0.246 |
| 256 | 4 | 36.811 | 23.630 | 7.959 | 0.779 | 0.249 |
| 512 | 2 | **37.983** | **24.260** | 7.896 | 0.770 | **0.253** |
| 1024 | 1 | 37.275 | 23.733 | 7.942 | 0.771 | 0.250 |

**Table S2:** Full sweep of protein-centric batch construction configurations for CARP 640M.

| P | M | EF@0.5 | EF@1.0 | EF@5.0 | AUCROC | BEDROC |
| --- | --- | --- | --- | --- | --- | --- |
| 1 | 1024 | 33.065 | 21.669 | 7.528 | 0.781 | 0.218 |
| 2 | 512 | 31.572 | 21.005 | 7.651 | 0.789 | 0.215 |
| 4 | 256 | 33.819 | 22.623 | 8.217 | **0.802** | 0.235 |
| 8 | 128 | 33.800 | 23.082 | 8.239 | 0.801 | 0.239 |
| 16 | 64 | 35.904 | 24.093 | **8.353** | 0.796 | 0.248 |
| 32 | 32 | 36.820 | 24.325 | 8.245 | 0.795 | 0.251 |
| 64 | 16 | 37.159 | 24.244 | 8.331 | 0.788 | 0.252 |
| 128 | 8 | 38.057 | 24.501 | 8.156 | 0.780 | 0.253 |
| 256 | 4 | 39.342 | **24.996** | 7.922 | 0.772 | 0.256 |
| 512 | 2 | 38.835 | 24.398 | 7.975 | 0.777 | 0.254 |
| 1024 | 1 | **39.568** | 24.871 | 8.112 | 0.773 | **0.258** |

**Table S3:** Full sweep of protein-centric batch construction configurations for ESM C.

| P | M | EF@0.5 | EF@1.0 | EF@5.0 | AUCROC | BEDROC |
| --- | --- | --- | --- | --- | --- | --- |
| 1 | 1024 | 29.918 | 19.954 | 7.251 | 0.787 | 0.203 |
| 2 | 512 | 29.827 | 20.211 | 7.436 | 0.778 | 0.207 |
| 4 | 256 | 28.491 | 20.084 | 7.776 | 0.792 | 0.208 |
| 8 | 128 | 29.767 | 20.485 | **7.869** | **0.796** | 0.214 |
| 16 | 64 | 30.473 | 20.729 | 7.781 | 0.791 | 0.217 |
| 32 | 32 | 30.333 | 20.583 | 7.658 | 0.784 | 0.214 |
| 64 | 16 | 29.057 | 19.857 | 7.520 | 0.783 | 0.209 |
| 128 | 8 | 29.661 | 20.232 | 7.507 | 0.778 | 0.212 |
| 256 | 4 | 32.079 | **21.221** | 7.404 | 0.769 | 0.220 |
| 512 | 2 | 31.571 | 21.051 | 7.365 | 0.769 | 0.218 |
| 1024 | 1 | **32.680** | 21.019 | 7.422 | 0.764 | **0.221** |

**Table S4:** Full sweep of protein-centric batch construction configurations for ESM1b.

| P | M | EF@0.5 | EF@1.0 | EF@5.0 | AUCROC | BEDROC |
| --- | --- | --- | --- | --- | --- | --- |
| 1 | 1024 | 36.455 | 24.062 | 7.875 | 0.792 | 0.242 |
| 2 | 512 | 37.574 | 24.247 | 8.204 | **0.797** | 0.249 |
| 4 | 256 | 38.652 | 25.136 | **8.444** | 0.794 | 0.257 |
| 8 | 128 | 38.757 | 24.966 | 8.427 | 0.792 | 0.260 |
| 16 | 64 | 37.875 | 24.812 | 8.280 | 0.790 | 0.256 |
| 32 | 32 | 38.519 | 24.723 | 8.118 | 0.780 | 0.256 |
| 64 | 16 | 37.721 | 24.630 | 8.200 | 0.779 | 0.255 |
| 128 | 8 | 39.637 | **25.532** | 8.226 | 0.774 | **0.263** |
| 256 | 4 | 39.951 | 25.512 | 8.125 | 0.764 | **0.263** |
| 512 | 2 | 39.845 | 24.974 | 7.904 | 0.760 | 0.259 |
| 1024 | 1 | **40.167** | 25.156 | 7.775 | 0.753 | 0.260 |

**Table S5:** Full sweep of protein-centric batch construction configurations for ESM2 T36.

| P | M | EF@0.5 | EF@1.0 | EF@5.0 | AUCROC | BEDROC |
| --- | --- | --- | --- | --- | --- | --- |
| 1 | 1024 | 35.968 | 23.481 | 7.948 | 0.785 | 0.238 |
| 2 | 512 | 36.467 | 24.179 | 8.217 | 0.792 | 0.245 |
| 4 | 256 | 38.245 | 25.156 | 8.517 | 0.795 | 0.258 |
| 8 | 128 | 37.570 | 24.774 | 8.437 | **0.798** | 0.255 |
| 16 | 64 | 37.715 | 24.739 | 8.670 | 0.795 | 0.258 |
| 32 | 32 | **39.615** | **25.520** | 8.626 | 0.797 | **0.267** |
| 64 | 16 | 39.458 | 25.499 | **8.682** | 0.791 | 0.266 |
| 128 | 8 | 39.154 | 24.967 | 8.392 | 0.783 | 0.262 |
| 256 | 4 | 39.724 | 25.059 | 8.232 | 0.775 | 0.262 |
| 512 | 2 | 38.989 | 24.603 | 8.023 | 0.771 | 0.257 |
| 1024 | 1 | 39.534 | 24.751 | 7.945 | 0.771 | 0.258 |

**Table S6:** Full sweep of protein-centric batch construction configurations for ProGen2.

| P | M | EF@0.5 | EF@1.0 | EF@5.0 | AUCROC | BEDROC |
| --- | --- | --- | --- | --- | --- | --- |
| 1 | 1024 | 21.891 | 14.048 | 5.452 | 0.741 | 0.144 |
| 2 | 512 | 21.381 | 14.724 | 6.091 | 0.753 | 0.153 |
| 4 | 256 | 22.888 | 16.150 | 6.524 | 0.768 | 0.167 |
| 8 | 128 | 24.091 | 16.935 | 6.886 | 0.777 | 0.178 |
| 16 | 64 | 28.013 | 19.046 | 7.171 | **0.784** | 0.199 |
| 32 | 32 | 28.684 | 19.714 | 7.291 | 0.776 | 0.206 |
| 64 | 16 | 31.261 | 20.639 | 7.255 | 0.769 | 0.214 |
| 128 | 8 | 32.392 | 21.194 | **7.515** | 0.777 | 0.221 |
| 256 | 4 | 33.023 | 21.405 | 7.301 | 0.768 | 0.222 |
| 512 | 2 | 34.794 | 22.364 | 7.273 | 0.760 | 0.231 |
| 1024 | 1 | **35.382** | **22.437** | 7.346 | 0.768 | **0.232** |

**Table S7:** Full sweep of protein-centric batch construction configurations for ProtBERT.

| P | M | EF@0.5 | EF@1.0 | EF@5.0 | AUCROC | BEDROC |
| --- | --- | --- | --- | --- | --- | --- |
| 1 | 1024 | 31.745 | 20.425 | 7.136 | 0.786 | 0.207 |
| 2 | 512 | 29.713 | 19.776 | 7.337 | 0.788 | 0.204 |
| 4 | 256 | 30.378 | 20.121 | **7.610** | **0.793** | 0.210 |
| 8 | 128 | 28.245 | 19.122 | 7.397 | 0.790 | 0.200 |
| 16 | 64 | 30.032 | 20.036 | 7.404 | 0.784 | 0.209 |
| 32 | 32 | 30.753 | 20.516 | 7.609 | 0.784 | 0.215 |
| 64 | 16 | 30.556 | 20.221 | 7.463 | 0.781 | 0.213 |
| 128 | 8 | 30.789 | 20.158 | 7.335 | 0.768 | 0.213 |
| 256 | 4 | 31.327 | 20.516 | 7.382 | 0.766 | 0.215 |
| 512 | 2 | 32.582 | 21.036 | 7.248 | 0.759 | 0.219 |
| 1024 | 1 | **33.332** | **21.420** | 7.226 | 0.759 | **0.222** |

**Table S8:** Full sweep of protein-centric batch construction configurations for ProtT5.

| P | M | EF@0.5 | EF@1.0 | EF@5.0 | AUCROC | BEDROC |
| --- | --- | --- | --- | --- | --- | --- |
| 1 | 1024 | **37.966** | **24.708** | 7.855 | 0.790 | **0.246** |
| 2 | 512 | 36.635 | 23.922 | 8.030 | 0.794 | 0.243 |
| 4 | 256 | 34.130 | 23.001 | 8.145 | **0.797** | 0.237 |
| 8 | 128 | 35.958 | 23.578 | 8.191 | 0.789 | 0.243 |
| 16 | 64 | 35.918 | 23.704 | **8.246** | 0.795 | 0.247 |
| 32 | 32 | 35.085 | 22.937 | 7.922 | 0.788 | 0.238 |
| 64 | 16 | 36.622 | 23.504 | 8.028 | 0.787 | **0.246** |
| 128 | 8 | 35.957 | 23.094 | 7.843 | 0.780 | 0.242 |
| 256 | 4 | 37.056 | 23.481 | 7.814 | 0.769 | 0.245 |
| 512 | 2 | 36.539 | 23.050 | 7.613 | 0.764 | 0.242 |
| 1024 | 1 | 36.949 | 23.274 | 7.594 | 0.757 | 0.243 |

## Encoder Selection

Table S9 presents the comparison of MolDeBERTa variants (MLM, MTR, and MLC) as molecular encoder for BindScreen. **Bold** indicates the best result for each metric within each variant.

**Table S9:**
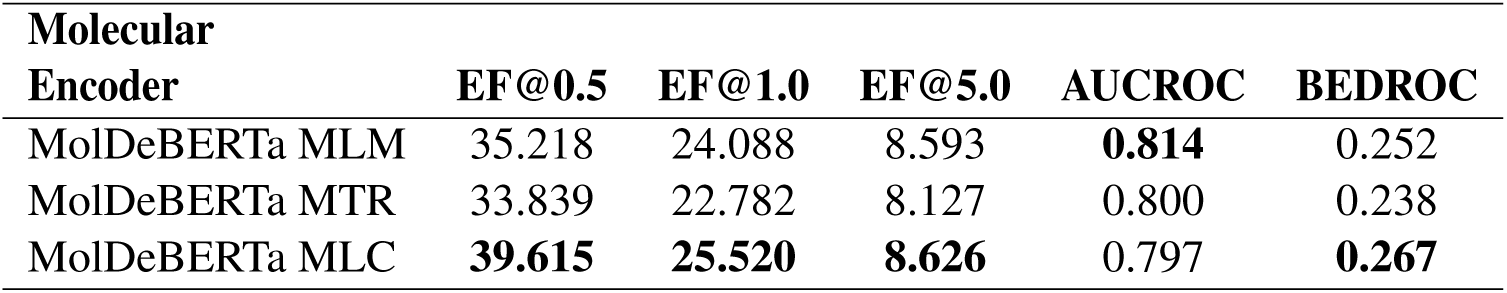
Comparison of MolDeBERTa variants as molecular encoder. **Bold** indicates the best result for each metric.

## Finetuning and Training Efficiency

Table S10 presents the comparison of finetuning strategies using ESM2 T36 and MolDeBERTa MLC. **Bold** indicates the best result for each metric within each variant.

**Table S10:**
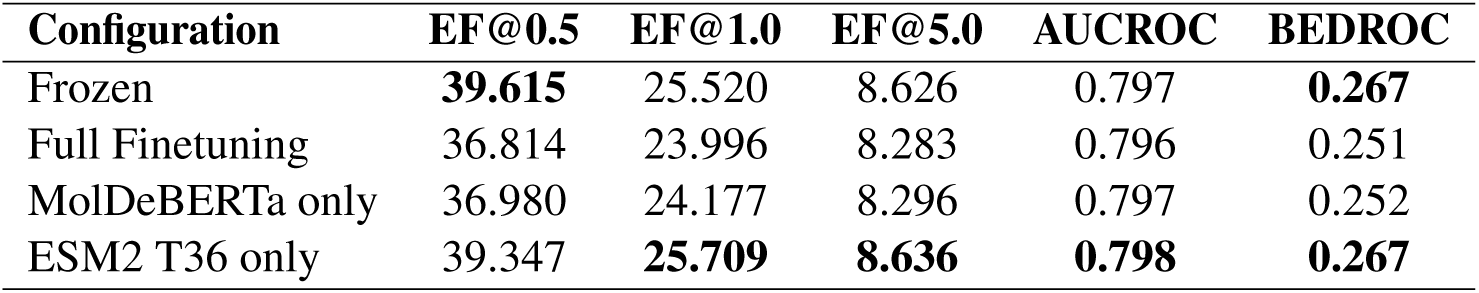
Comparison of finetuning strategies using ESM2 T36 and MolDeBERTa MLC. **Bold** indicates the best result for each metric.

| Configuration | EF@0.5 | EF@1.0 | EF@5.0 | AUCROC | BEDROC |
| --- | --- | --- | --- | --- | --- |
| Frozen | **39.615** | 25.520 | 8.626 | 0.797 | **0.267** |
| Full Finetuning | 36.814 | 23.996 | 8.283 | 0.796 | 0.251 |
| MolDeBERTa only | 36.980 | 24.177 | 8.296 | 0.797 | 0.252 |
| ESM2 T36 only | 39.347 | **25.709** | **8.636** | **0.798** | **0.267** |

## Footnotes

1 https://github.com/KSUN63/DeepDTA-Pytorch

2 https://github.com/zhaoqichang/HpyerAttentionDTI

3 https://github.com/kexinhuang12345/MolTrans

